# Cullin3-RING ubiquitin ligases are intimately linked to the unfolded protein response of the endoplasmic reticulum

**DOI:** 10.1101/428136

**Authors:** Kyungho Kim, Tamara Moretti, Sujin Park, Jinoh Kim

## Abstract

CUL3-RING ubiquitin ligases (CRL3s) are involved in various cellular processes through different Bric a brac, Tramtrack and Broad-Complex (BTB)-domain proteins. KLHL12, a BTB-domain protein, appears to play an essential role in export of large cargo molecules like procollagen from the endoplasmic reticulum (ER). It has been suggested that CRL3^KLHL12^ mono-ubiquitinates SEC31 and mono-ubiquitinated SEC31 increases the dimension of a COPII coat to accommodate the large cargo molecules. As we examined this model, we found that functional CRL3^KLHL12^ was indeed critical for the assembly of large COPII structures. Interestingly, we noticed that CRL3^KLHL12^ influences collagen synthesis in human skin fibroblasts (HSFs). Our results also suggest that there is a CRL3^KLHL12^–independent collagen secretion route in HSFs. In addition, we found that CRL3^KLHL12^ strongly influences levels of sensors of the unfolded protein response (UPR). Different cell lines reacted differently to CUL3 depletion with respect to UPR regulation. This cell line-dependency appears to rely on a cell line-specific BTB-domain protein(s). Consistent with this idea, depletion of a muscle-specific BTB-domain protein KLHL41 recapitulated the effects of CUL3 depletion in C2C12 myotubes in UPR regulation. Based on these results we propose that CRL3^KLHL12^ and CRL3^KLHL41^ are regulators of the UPR.

## INTRODUCTION

CUL3-RING ubiquitin (Ub) ligases (CRL3s) are the major E3 ligase family in eukaryotes [1]. A CRL is composed of a catalytic module and a substrate recognition module [1]. In CRL3s, CUL3 and RBX1 serve as the catalytic module and a Bric a brac, Tramtrack and Broad-Complex (BTB) domain protein serves as the substrate recognition module [2-4]. There are 183 BTB domain genes in the human genome [5], indicating that CUL3 is involved in various cellular processes through the BTB domain proteins. Indeed, recent studies have shown that CRL3s are important regulators of different cellular and developmental processes such as mitosis, cytokinesis, cell death, gene expression control, Hedgehog signaling, WNT signaling and so on where different BTB domain proteins are involved (see Genschik et al. for review [6]). Defects of CRL3s have been implicated in metabolic diseases, dystrophies, and cancers [6].

KLHL12 is a BTB domain protein and CRL3^KLHL12^ appears to play an important role in export of procollagen from the ER by ubiquitinating SEC31, a component of a COPII coat [7, 8]. The COPII coat consisting of SAR1, SEC23/SEC24 complex, and SEC13/SEC31 complex is able to generate transport vesicles at the ER exit sites that can package most soluble and membrane proteins destined to cellular locations. Collagens are the major component of the extracellular matrix (ECM). Type I and II procollagens form about 300 nm-long rigid triple helices [9, 10] and require COPII proteins for export from the ER [11-17]. Because COPII proteins typically generate vesicles limited to 60-80 nm in diameter, a mechanism must exist that allows COPII coat proteins to accommodate such large cargo molecules. Alternatively, a COPII-dependent short-loop pathway that does not involve large vesicles account for transport of collagen from the ER to the Golgi [18].

When KLHL12 was overexpressed, larger COPII-coated vesicles (>500 nm) were observed [7] and collagens were found in large structures coated with COPII and KLHL12 [19]. Furthermore, large COPII structures were generated even in the presence of lysine-free ubiquitin which prevents poly-ubiquitination of SEC31 [7]. Based on these results, mono-ubiquitinated SEC31 molecules contribute to enlargement of COPII vesicles and packaging of procollagen into large vesicles.

Despite the evidence for the essential role of KLHL12 in collagen secretion, its effect on collagen secretion is relatively modest. For example, depletion of COPII or TANGO-1 displays a strong effect on collagen deposition in the ECM or medium [20-22]. However, depletion of KLHL12 requires an integrin signaling inhibitor to exhibit a collagen phenotype although depletion of CUL3 alone disrupted a collagen deposition in the ECM in mouse embryonic stem cells [7]. In addition, KLHL12 is downregulated upon differentiation [7], posing a question if KLHL12 is essential for collagen secretion under every circumstances. As we investigated the proposed role of CUL3-KLHL12 in collagen secretion, we found out that KLHL12 is not essential for collagen secretion in human skin fibroblasts and we discovered new functions of CRL3s in collagen synthesis as well as in the unfolded protein response (UPR) of the ER.

## MATERIALS AND METHODS

### Plasmid and siRNA

The pcDNA5/FRT/TO KLHL12-FLAG construct was a gift of Michael Rape’s laboratory (University of California, Berkeley, CA, USA). The KLHL12 Mut A (ELSE65-68AAAA), Mut B (CLLQLK111-116AAAAAA), and Mut C (FAETHNC141-147AAAAAAA) were generated by site-directed mutagenesis (Fig. S1 and Table S1). pCMV6-Entry (PS100001), human pCMV6-KLHL40-Myc-DDK (RC213832), and human pCMV6-KLHL41-Myc-DDK (RC200295) were purchased from Origene (Rockville, MD, USA). The control (D-01910-10-50), human–specific CUL3 (M-010224-02), and KLHL12 siRNAs (M-015890-02), mouse-specific CUL3 (M-062951-01-0005), KLHL40 (M-016211-02-0005), and mouse-specific KLHL41 (M-045964-01-0005) were purchased from Dharmacon (Lafayette, CO, USA). Sequences of these siRNAs are shown as follows. Control (pool of 4) siRNAs: 5’-UAGCGACUAAACACAUCAA-3’, 5’-**UAAGGCUAUGAAGAGAUAC**-3’, 5’-AUGUAUUGGCCUGUAUUAG-3’, 5’-AUGAACGUGAAUUGCUCAA-3’. Human CUL3 (pool of 4) siRNAs: 5’-**CCGAACAUCUCAUAAAUAA**-3’, 5’-**GAGAAGAUGUACUAAAUUC**-3’, 5’-**GAGAUCAAGUUGUACGUUA**-3’, 5’-GCGGAAAGGAGAAGUCGUA-3’. Mouse CUL3 (pool of 4) siRNAs: 5’-**GAAAGUAGAAGCUAGGAUU**-3’, 5’-**CGGCAAACUUUAUUGGAUA**-3’, 5’-**GGUGAUGAUUAGAGACAUA**-3’, 5’-**CCAAGUCCAGUUGUUAUUA**-3’. Human KLHL12 (pool of 4) siRNAs : 5’-GGAAGGUGCCGGACUCGUA-3’, 5’-GCGCUAUGAUCCAAACAUU-3’, 5’-GAAGAAUCCUUGCCUAACC-3’, 5’-UAAUCAAGUGCGACGAAAU-3’. Mouse KLHL40 (pool of 4) siRNAs: 5’-**GCGCAUACUUCCUGCAGUU**-3’, 5’-GAACAAGAUGUGCGUCUAU-3’, 5’-**CUACGUAAUUGGCGGCAAA**-3’, 5’-**AUGAGGAGGCCGAACGUAU**-3’. Mouse KLHL41 (pool of 4) siRNAs: 5’-GGGAAGUGAUGACGGAAUU-3’, 5’-GUUCAUCAUUGCACCCUA-3’, 5’-GGCAUGGAAUGUUUGUUAA-3’, 5’-CCGGUUACCUGAACGAUAU-3’

### Antibodies and chemicals

Rabbit anti-PC1 antibodies were a gift from L. Fisher (LF-68, 1:8000, National Institute of Dental and Craniofacial Research, Bethesda, MD, USA)[23]. Rabbit anti-ribophorin I antibody (1:5,000) was a gift from Schekman laboratory (University of California, Berkeley, CA, USA). Additional antibodies we used include rabbit anti-CUL3 (A301-109A, 1:1000, Bethyl Laboratories, Universal Biologicals, Cambridge, UK), mouse FK2 anti-Ub (BML-PW8810, 1: 1000, Enzo Life Sciences, Farmingdale, NY, USA), anti-FLAG (F3165, F7425, 1:1500, Sigma-Aldrich, St. Louis, MO, USA), mouse anti-KLHL12 (#30058, 1:1000, ProMab Biotechnology, Richmond, CA,USA), rabbit anti-KLHL40 (#HPA024463, 1:1000, Sigma-Aldrich, St. Louis, MO, USA), rabbit anti-KLHL41 (#AV38732, 1:1000, Sigma-Aldrich, St. Louis, MO, USA), mouse and rabbit anti-SEC31A (#612350, BD Biosciences, San Diego, CA, A302-336A, Bethyl Laboratories, Universal Biologicals, Cambridge, UK), rabbit anti-IRE1α (#3294), PERK (#5683), p-eIF2α (Ser51, #9721), total-eIF2α (#9722), XBP-1s (#12782) (1:1000 final dilution; all from Cell signaling Technology, Danvers, MA, USA), monoclonal anti-ATF6α antibody (73-500, 1:1200, B-Bridge International, Inc., Santa Clara, CA, USA), monoclonal anti-α-tubulin (TU-02, sc-8035, 1:2000, Santa Cruz Biotechnology, Dallas, TX, USA). Hygromycin B (10687-010) and blasticidin S (R210-01) were purchased from Invitrogen (Carlsbad, CA, USA). Cycloheximide (CHX, C4859), dimethyl sulfoxide (DMSO, D4540,) and 1, 4-dithiothreitol (DTT, D0632) were purchased from Sigma-Aldrich (St. Louis, MO, USA). Protease inhibitor cocktail (Mannheim, Germany), MLN4924 (15217, Cayman Chemical, Ann Arbor, MI, USA), MG132 (474790, Calbiochem, San Diego, CA, USA), brefeldin A (BFA, #9972, Cell signaling Technology, Danvers, MA, USA) were also used.

### Cell culture and transfection

Normal human skin fibroblasts (CRL-2091), HeLa cells (HeLa, CCL-2), human lung fibroblasts (IMR-90, CCL-186), and mouse C2C12 myoblasts (CRL1772) were purchased from the American Type Culture Collection (ATCC, Manassas, VA, USA). Cells were maintained in low-glucose Dulbecco’s Modified Eagle Medium (DMEM) with 10% fetal bovine serum (FBS) and 1% penicillin–streptomycin. Flp_In T-REx^™^ 293 host cell line containing a stably integrated FRT site and tetracycline (Tet) repressor was grown in high-glucose DMEM, supplemented with 10% FBS, blasticidin and hygromycin B as instructed by the manufacturer (Gibco/Thermo Fisher, Waltham, MA, USA) and gene expression was induced by 1 μg/ml doxycycline. Cells were maintained in high-glucose Dulbecco’s Modified Eagle Medium (DMEM) with 10% fetal bovine serum (FBS) and 1% penicillin–streptomycin for proliferation (Gibco/Thermo Fisher, Waltham, MA, USA). Differentiation C2C12 myoblasts into myotubes was initiated by incubating the cells in high-glucose Dulbecco’s Modified Eagle Medium (DMEM) with 2 % Horse serum (HS) (Thermo Fisher, Waltham, MA, USA) and 1% penicillin-streptomycin [24]. Cell lines were maintained in a water jacketed incubator at 37°C with 5% CO_2_ enrichment. Lipofectamine LTX 3000 was used for plasmid DNAs transfection and Lipofectamine RNAiMAX for siRNAs transfection according to the manufacturer’s protocols unless specified otherwise. All of the cell culture and transfection reagents were purchased from Invitrogen (Carlsbad, CA, USA).

### Establishing stable cell lines

Flp-In™ T-REx™ 293 cells stably expressing Tet-inducible KLHL12 WT, Mut A, and Mut C were generated according to the manufacturer’s instructions (Flp-In™ System, Invitrogen, Carlsbad, CA, USA). Briefly, the Flp-In™ T-REx™ 293 Cells (R780-07) were plated onto 6-well plates and incubated for 24 h. The following day, the cells were transfected using 5 μl Lipofectamine LTX with 0.5 μg of a KLHL12 construct and 1.0 μg of pOG44 (V6005-20). After 48 h post-transfection, the cells were replated on 24-well plates and subjected to selection with hygromycin B (100 μg/ml) and blasticidin S (10 μg/ml) for up to 4 weeks. *KLHL12* expression was induced by 1 μg/ml doxycycline.

### Co-immunoprecipitation and Immunoblotting

For co-immunoprecipiation, cells were collected by centrifugation and lysed by douncing 30 times using buffer A (0.1% NP-40, 2.6 mM KCl, 1.5 mM KH_2_PO_4_, 140 mM NaCl, 8 mM Na_2_HPO_4_-7H_2_O, and 1X protease inhibitor cocktail) on ice. After centrifugation for 15 min at 14,000 rpm at 4°C, cleared lysates were incubated with indicated antibodies overnight in the cold room, followed by incubation with protein A or protein G Sepharose beads (GE Healthcare, Milwaukee, WI, USA) for 3 h with gentle rocking in the cold room. The beads were harvested by centrifugation and washed three times with buffer A. After adding the SDS-sample buffer and heating for 5 min at 95 °C, proteins were resolved with SDA-PAGE and processed for immunoblotting.

For routine immunoblotting analysis, cells were lysed with buffer B (20 mM Tris-HCl, pH 7.5, 150 mM NaCl, 1 mM Na_2_EDTA, 1 mM EGTA, 1% NP-40, 1% sodium deoxycholate, 2.5 mM sodium pyrophosphate, 1 mM β-glycerophosphate, 1 mM Na_3_VO_4_, 1X complete protease inhibitor cocktail) on ice for 15 minutes. Cell lysates were cleared by centrifugation at 14,000 rpm for 15 minutes at 4°C. Cleared lysates were quantified using Pierce BCA Protein Assay Kit (Thermo Fisher Scientific, Waltham, MA, USA). Equal amounts of proteins were resolved on 6 or 10% SDS-PAGE, transferred to PVDF membrane (Millipore, Bedford, MA, USA), probed with primary antibodies, subsequently with horseradish peroxidase-conjugated secondary antibodies, and visualized with ECL Prime (GE Healthcare, Pittsburgh, PA, USA). Band intensity was quantified using the ImageJ software. Protein levels were normalized to ribophorin I or α-tubulin and are presented as mean ± standard deviation.

### Ubiquitination assay

The ubiquitination assay was performed as previously described [25]. Briefly, Tet-inducible KLHL12-FLAG-expressing 293 cells were treated with MG132 for 24 h prior to harvest. To assess endogenous ubiquitination of KLHL12, cells were harvested in PBS containing 5 mM NEM and lysed in 1% SDS by boiling for 10 min. Cell lysates were diluted to 0.1% SDS by adding lysis buffer (6 M guanidinium-HCl buffer (pH 8.0)) containing protease inhibitors and 5 mM NEM, and immunoprecipitated with anti-FLAG antibody followed by immunoblot.

### Immunofluorescent microscopy

Cells were plated in 6-well plate with acid-washed glass coverslips. After 24h, cells were fixed with 4% paraformaldehyde (PFA) in DPBS for 30 min, washed five times with PBS and permeabilized with 0.1% Triton X-100 in DPBS for 15 min at room temperature. Primary antibodies were used in 0.5% BSA in DPBS. Cells were incubated with blocking buffer (0.5% BSA in PBS) for 30 min at RT followed by 1 h incubation at RT with rabbit anti-FLAG antibody (1:500, Sigma-Aldrich, St. Louis, MO, USA) and mouse anti-SEC31A antibody (1:500, BD Biosciences, NJ, USA) and then secondary antibodies such as Alexa Fluor 488 goat anti-rabbit IgG and Alexa Fluor 594 goat anti-mouse IgG (1:500, Molecular Probes, Invitrogen, Carlsbad, CA, USA). Antibody incubations were followed by five PBS washes. Images were obtained using a confocal laser microscopy (Nikon C1, Nikon, Tokyo, Japan), and merges of images were processed using Nikon EZ-C1 viewer software.

### Real-time quantitative PCR with reverse transcription

RNA was obtained from fibroblast cells using Trizol reagent (Invitrogen, Carlsbad, CA, USA) according to the manufacturer’s instructions. Complementary DNAs (cDNAs) were synthesized from total RNA using Superscript II First-strand Synthesis kit (Invitrogen, Carlsbad, CA, USA) and quantified by real-time PCR using the IQ SYBR green supermix kit (Bio-Rad Laboratories, Hercules, CA, USA) and the following primers: human *COL1A1*, 5’-GTGTTGTGCGSTGSCG-3’ and 5’-TCGGTGGGTGACTCT-3’; human *CUL3*, 5’-CTGGTGTATCTTTAGGTGGTG-3’ and 5’-GTGCTGGTGGGATGTTG-3’; human *GAPDH*, 5’-CTCCTCCACCTTTGACGC-3’ and 5’-CCACCACCCTGTTGCTGT-3’. Gene expression levels were normalized to *GAPDH* expression levels.

### Statistical analysis

Statistical analyses were performed using GraphPad Prism (GraphPad Software Inc., La Jolla, CA, USA). The results of multiple experiments were presented with meanL±Lstandard deviation (SD). P values of less than 0.05 were considered statistically significant.

## RESULTS

### MLN4924 is a potent inhibitor of CRL3s

The catalytic activity of CRL3^KLHL12^ has been shown to be critical for the assembly of large COPII vesicles and collagen export from the ER [7]. To modulate the activity of CRL3^KLHL12^, we searched for small molecules that can inhibit the Ub ligase activity of CRL3^KLHL12^. Cullin itself is not a catalytic enzyme, but a scaffold protein. The Ub ligase activity of CRL3 is activated by neddylation of CUL3 as CUL3 neddylation induces a drastic conformational change near the RBX1 binding region [26]. This conformational change results in juxtaposition of E2-Ub to a substrate [27, 28]. MLN4924 is a potent inhibitor of NEDD8-activating enzyme and strongly inhibits the catalytic activity of CRLs [29]. We used this small molecule to survey its effect on the catalytic activity of CRL3 and the interactions among the components of the CRL3^KLHL12^ complex. For this purpose, we established a doxycycline-inducible HEK cell line stably expressing FLAG tagged KLHL12 (KLHL12-FLAG). Interestingly, MLN4924 treatment not only prevented CUL3 neddylation (Fig. 1A), but also stabilized KLHL12 (Fig. 1A, Input; Fig. S3, WT). This result suggests that turnover of KLHL12 is regulated by neddylation of CRLs. When we immunoprecipitated KLHL12, similar levels of SEC31 or CUL3 were co-precipitated regardless of inhibitor treatment (Fig. 1A, IP). Although apparent levels of SEC31 and CUL3 bound to KLHL12 were not affected by MLN4924 treatment, because more KLHL12 molecules were present due to stabilization there was a decrease in the relative amounts of SEC31 bound to KLHL12. Thus, neddylation of CUL3 seems to help form CRL3^KLHL12^-SEC31 complex more effectively.

**Figure 1.**
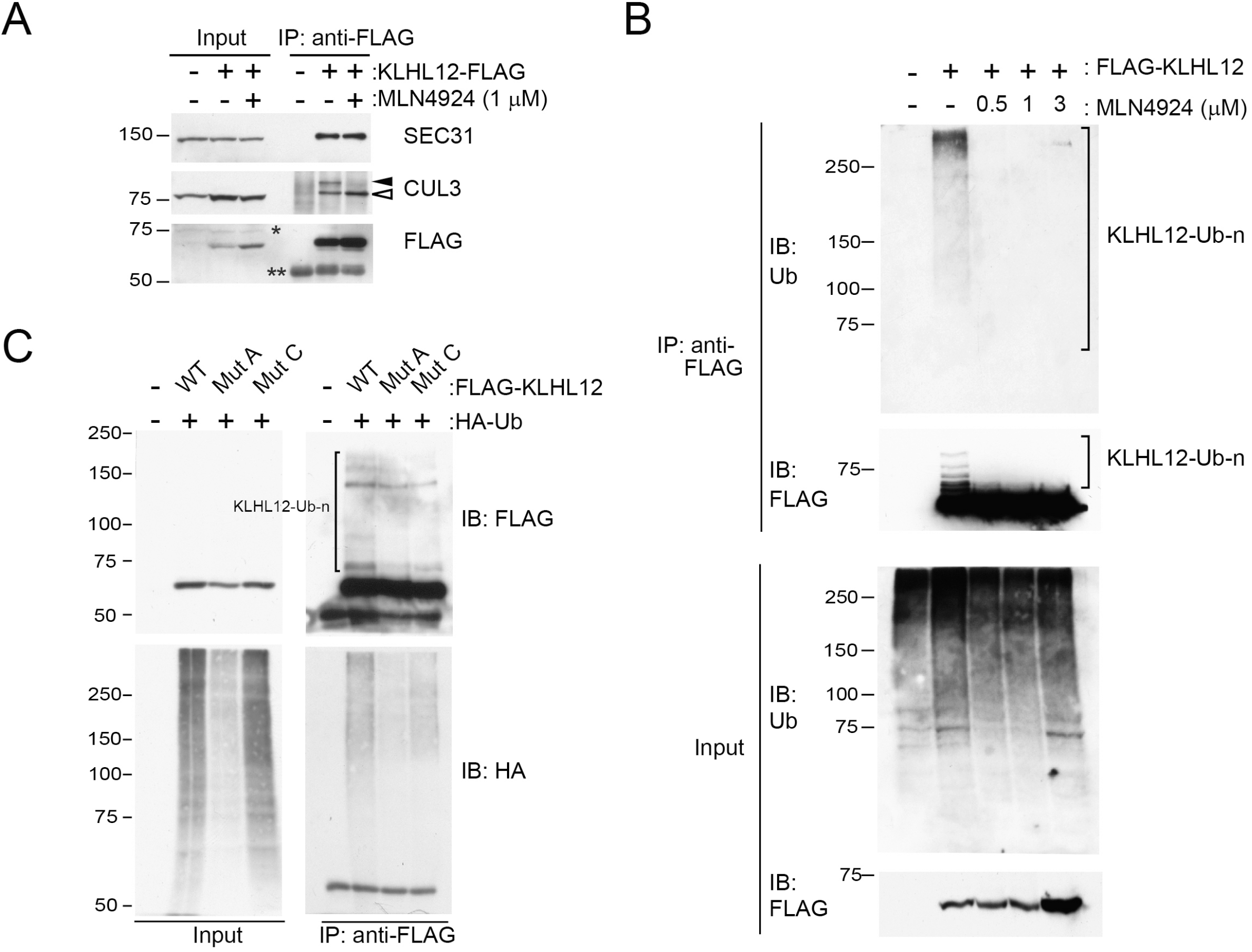
MLN4924, an inhibitor of NEDD8-activating enzyme, inhibits the activity of CRL3^KLHL12^. (A and B) 293 cells stably expressing KLHL12-FLAG were incubated in the presence or absence of MLN4924. Expression of KLHL12-FLAG was induced with doxycycline. Cellular lysates were processed for immunoprecipitation using an anti-FLAG antibody. Closed arrowhead, neddylated CUL3; open arrowhead, unneddylated CUL3. *Nonspecific band. **Immunoglobulins. (C) An HA-tagged ubiquitin construct was expressed in HEK cells stably expressing wildtype (WT) or a mutant KLHL12 (Mut A or Mut C). Cellular lysates were processed for immunoprecipitation using an anti-FLAG antibody. The expression of HA-Ub was low in the cells expressing Mut A.

We then asked whether MLN4924 inhibits the catalytic activity of CRL3s. Unfortunately, however, ubiquitinated forms of endogenous SEC31 were not detectable under our experimental conditions. To circumvent this problem, we took advantage of the observation that turnover of KLHL12 is stabilized by MLN4924 (Fig. 1A). It has been reported that an overactive form of CUL3 degrades KLHL3 by facilitating KLHL3 poly-ubiquitination and MLN4924 stabilizes KLHL3 by inhibiting CUL3-dependent KLHL3 poly-ubiquitination [30]. In analogy, KLHL12 may be ubiquitinated by CUL3. If this is the case, MLN4924 would inhibit KLHL12 ubiquitination. Indeed, MLN4924 treatments inhibited ubiquitination of KLHL12 (Fig. 1B), consistent with the idea that CUL3 ubiquitinates KLHL12. To corroborate this idea, we generated KLHL12 constructs with mutations at the KLHL12-CUL3 interface (Fig. S1 and Table S1, Mut B was not expressed for unknown reasons). Mut A or Mut C showed defects in CUL3 binding (Fig. S2A, B). In addition, they were ubiquitinated with reduced efficiency (Fig. 1C). Our results indicate that KLHL12 ubiquitination is CUL3-dependent. Taken together, our data suggest that MLN4924 is a potent inhibitor of CRL3^KLHL12^.

Interestingly, we noticed a strong reduction of neddylated (ned)-CUL3 bound to Mut A or Mut C when evaluated by immunoprecipitation (Fig. S2B). This was not due to lack of ned-CUL3 as we observed ned-CUL3 in total cell lysates prepared from cells stably expressing the KLHL12 constructs (Fig. S3). Thus, our data suggest that Mut A and Mut C do not form stable ned-CUL3-KLHL12 complexes. Mut A and Mut C also showed defects in KLHL12-SEC31 interaction (Fig. S2C, D). Clearly, the KLHL12-CUL3 interface is important not only for KLHL12-CUL3 interaction, but also for KLHL12-SEC31 interaction. Because ned-CUL3 species were deficient in mutant KLHL12-CUL3 complexes, this lack of ned-CUL3 in the complex likely contributed to the defective SEC31 binding. A similar finding was observed with KLHL12-SEC31 interaction in the presence of MLN4924 (Fig. 1). Note that although apparent levels of SEC31 and CUL3 bound to KLHL12 were not affected by MLN4924 treatment, there was a decrease in the relative amounts of SEC31 bound to KLHL12 due to KLHL12 stabilization.

### Formation of large COPII structures relies on functional CRL3^KLHL12^

Because MLN4924 prevents neddylation of CUL3, an essential step for CRL activation, we asked whether this inhibitor blocks the formation of large COPII-coated structures. To test this possibility, we treated the 293 cells stably expressing KLHL12 with MLN4924 and monitored COPII-KLHL12-coated structures via immunofluorescent microscopy. As previously reported, overexpression of KLHL12 induced formation of large COPII-KLHL12-coated structures (Fig. 2A, DMSO panel). In the presence of MNL4924, we observed aberrant COPII-KLHL12 structures with reduced SEC31 signals (Fig. 2A, MLN4924 panels), suggesting that the catalytic activity of CRL3^KLHL12^ is necessary for formation of large COPII coats.

**Figure 2.**
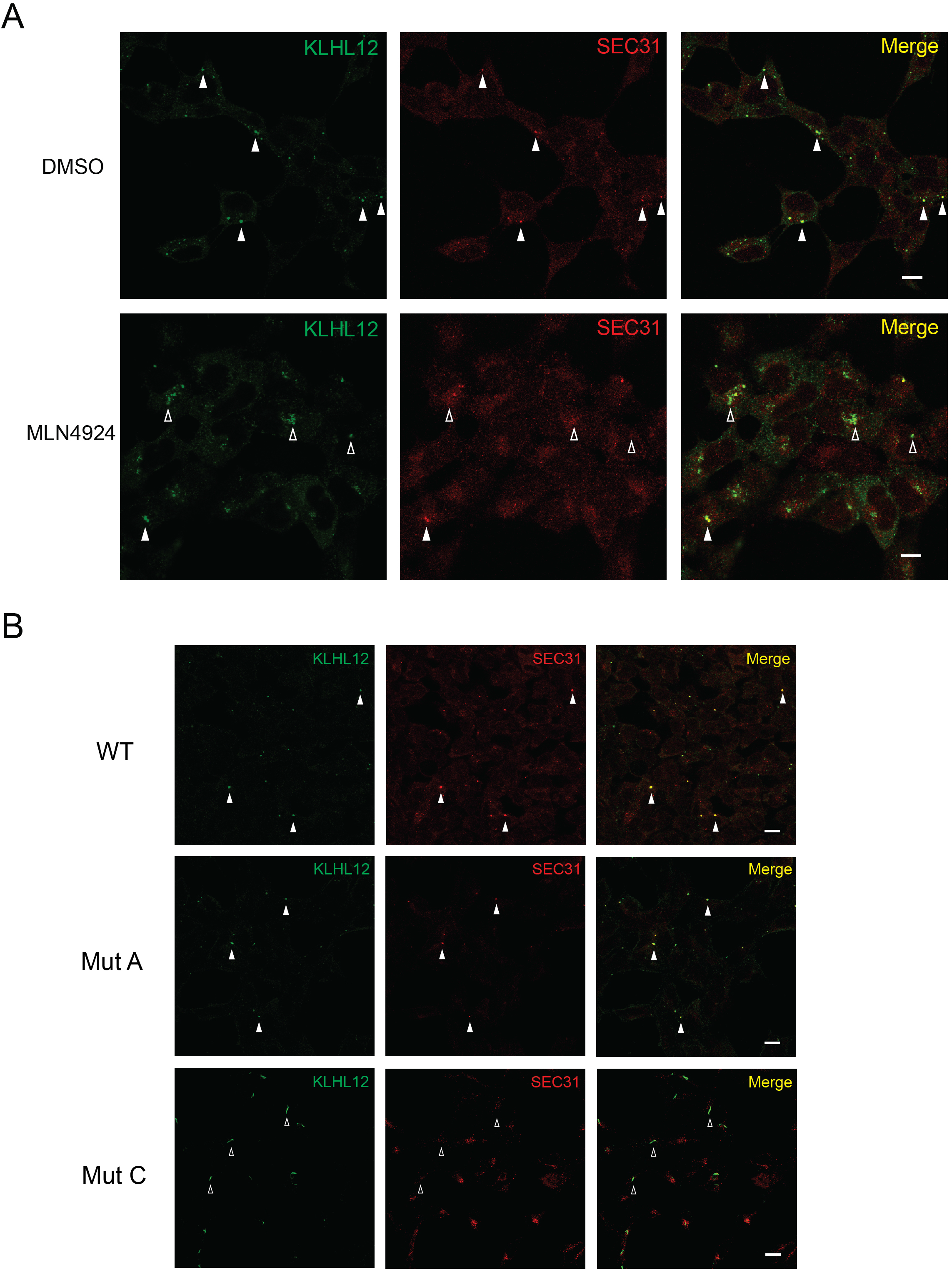
Aberrant large COPII structures with compromised CRL3^KLHL12^. (A) 293 cells stably expressing WT KLHL12-FLAG were incubated in the presence (1μM) or absence (DMSO) of MLN4924 and processed for immunofluorescent labeling. Labeled cells were visualized by standard confocal microscopy. Aberrant large COPII-KLHL12 structures were formed upon the MLN4924 treatment. (B) 293 cells stably expressing the indicated construct were processed for immunofluorescent labeling. Labeled cells were visualized by standard confocal microscopy. Large structures that are co-stained with KLHL12 and SEC31A were marked with closed arrowheads. Those that are deficient with SEC31A labeling were marked with open arrowheads.

We also monitored large COPII structures induced by expression of Mut A or Mut C (Fig. 2B). Interestingly, Mut A induced formation of apparently normal COPII-KLHL12 structures except that intensities of SEC31 fluorescence were slightly weaker than those of WT (Fig. 2B). Mut C induced formation of elongated COPII-KLHL12 structures with strongly reduced SEC31 fluorescence. These results reflect reduced interactions between the mutant KLHL12s and SEC31 (Fig. S2C, D). Taken together with the MLN4924 results, our data suggest that a proper KLHL12-CUL3 interaction and CUL3 neddylation are important for formation of large COPII-KLHL12 structures.

### MLN4924 reduces the intracellular levels, but not the secretion of collagen

As MLN4924 treatment induced the formation of aberrant structures, we expected that this inhibitor would reduce collagen secretion. We used human dermal fibroblasts (HSFs) which secrete type I collagen robustly because the 293 cells do not express type I collagen (Fig. S4). Intracellular collagen molecules were efficiently depleted within 3h in HSFs when new collagen synthesis was blocked (Fig. 3A). MLN4924 treatment inactivated CRL3s as indicated by the absence of ned-CUL3 (Fig. 3B, see the CUL3 panel). Unexpectedly, however, secretion of collagen was not affected by MLN4924 treatments (Fig. 3B and C), but the levels of intracellular collagen were reduced by the inhibitor treatments (Fig. 3B and D). Prolonged inhibitor treatments (> 24h) blocked additional deposition of collagen in the medium (Fig. S5, see 48h and 72h). However, this was due to complete depletion of collagen in the cells (see lysate). These data suggest that CRLs primarily regulate cellular collagen levels. While our data gave us an interesting clue toward a new role of CUL3 in collagen, we recognized that MLN4924 inhibits various cellular processes involving NEDD8-activating enzyme [29].

**Figure 3.**
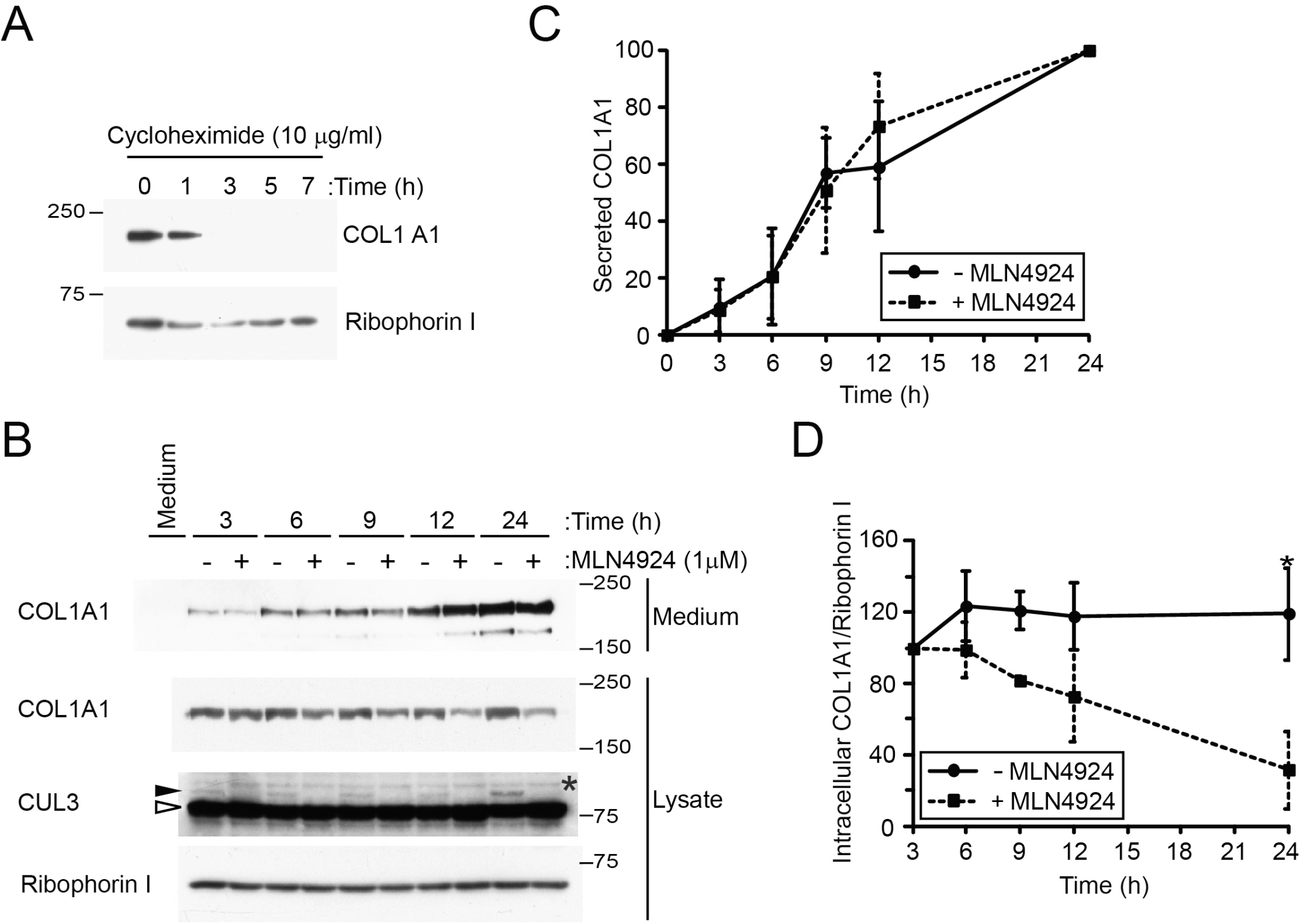
Regulation of cellular COL1A1 levels by MLN4924. (A) Rapid depletion of intracellular collagen was observed in human skin fibroblasts (HSFs) after cycloheximide treatment. (B) HSFs were plated and incubated for 24h. Afterwards, the culture medium was replaced with fresh medium with or without MLN4924 (1μM). Conditioned media and cells were collected at indicated times and processed for immunoblotting. The amount of a medium loaded into the gel was normalized to the amount of total proteins in a cell lysate. Riboporin I was probed as a loading control. Medium, unconditioned medium; closed arrowhead, ned-CUL3; open arrowhead, unneddylated-CUL3. (C, D) Quantification of levels of secreted (C) and intracellular (D) collagen. Protein bands were quantified using ImageJ. Student’s t-test: P*<0.05, n=3. (E, F) Depletion of KLHL12 (E) or CUL3 (F) by siRNA. Asterisks represent nonspecific proteins. Occasionally, a proteolytic processing of collagen occurs in the secreted collagen, resulting in a fast migrating species (open arrowheads). Note that the reduction in the levels of KLHL12 or CUL3 already reached the maximum at the lowest siRNA concentration used.

### CUL3 is a regulator of COL1A1 expression

To test if CUL3 is indeed responsible for regulation of collagen levels we used an approach of RNA interference (RNAi) with small interfering RNAs (siRNAs) (Fig. 4A, B). Initially, we performed RNAi in the absence of fetal bovine serum (FBS) during a transfection step (16 h) where cells are exposed to transfection reagents to maximize the transfection efficiency. Subsequently, we realized that inclusion of FBS during this step yielded more robust results. When the serum was present during the transfection step, intracellular collagen levels increased (Fig. 4A, C)(compare lane 1 with lane 3; lane 2 with lane 4). This result is consistent with previous reports that the serum enhances synthesis of type I collagen in HSFs [31, 32].

**Figure 4.**
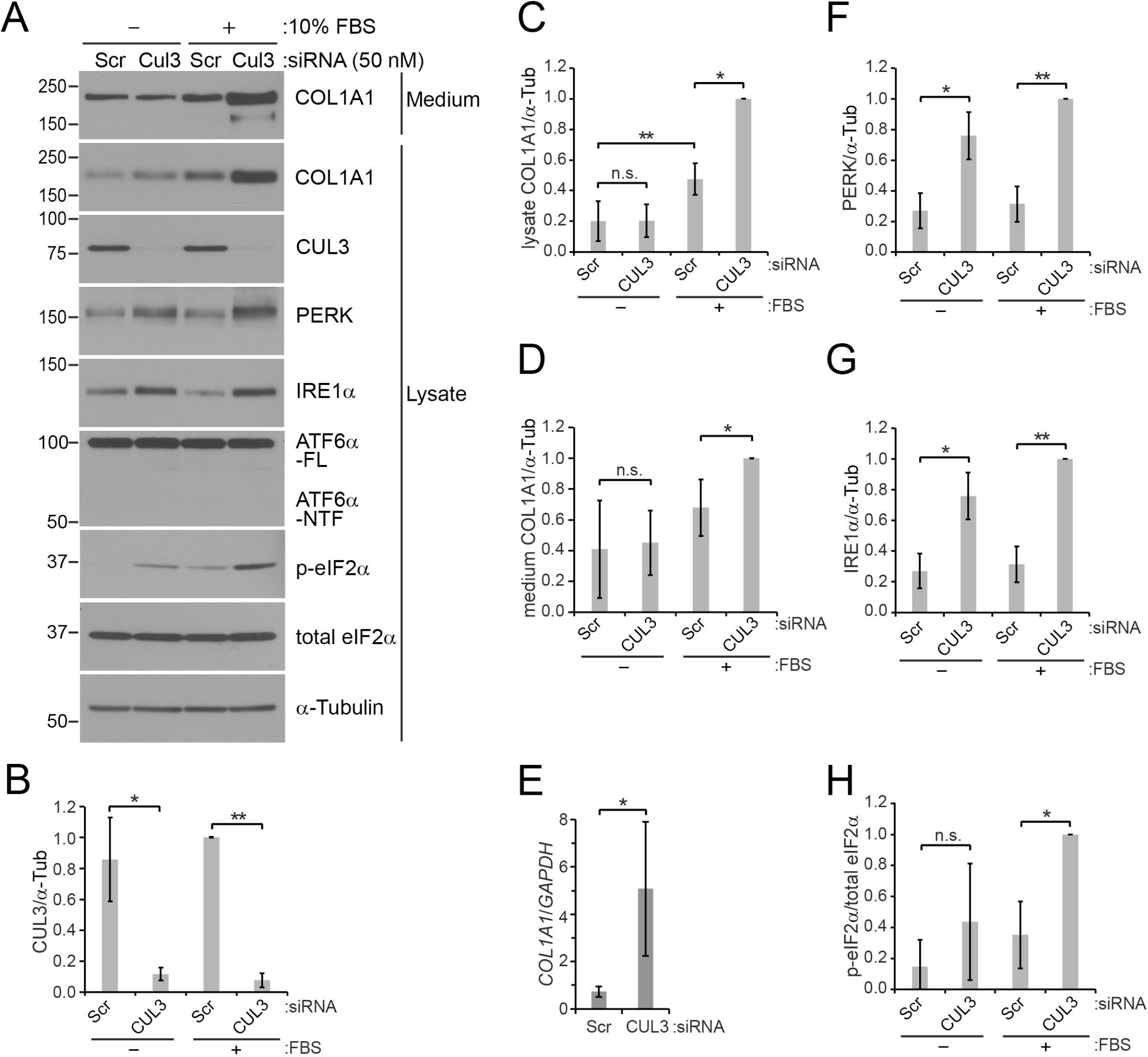
Regulation of COL1A1, PERK, and IRE1α levels by CUL3. (A-G) HSFs were transfected with scrambled (Scr) or CUL3 siRNAs. When the cells were incubated with siRNAs and transfection reagents (about 16 h), 10% FBS was absent or present as indicated. The culture medium was then replaced with a fresh complete medium containing 10% FBS. The cells and the conditioned media were collected after additional 48 h of incubation. (A) A representative immunoblot images. (B-D, F-H) Immunoblots images of 4-5 independent experiments were quantified. Statistical analyses were performed with Student’s t-test. Error bars represent standard deviations. (B) Comparison of CUL3 levels normalized to α-tubulin. P*<0.0005; P**<0.0001, n=5. (C) Comparison of intracellular COL1A1 levels. P*<0.0001; P**<0.01, n=5. n.s., non-significant. (D) Comparison of secreted COL1A1 levels. P*<0.005, n=5. n.s., non-significant. (E) Levels of *COL1A1* mRNAs were measured with RT-qPCR and normalized to levels of *GAPDH* mRNAs. P*<0.005, n=7. (F) Comparison of PERK levels. P*<0.0005; P**<0.0001, n=5. (G) Comparison of IRE1α levels. P*<0.0005; P**<0.0001, n=5. (H) Comparison of phosphorylated (p)-eIF2α levels normalized to total levels of eIF2α. P*=0.001, n=4. n.s., non-significant.

Depletion of CUL3 enhanced the levels of intracellular and secreted collagen especially when the serum was present (Fig. 4A, C, D)(compare lane 3 with lane 4). This was unexpected because the MLN4924 treatment reduced collagen levels. However, it should be noted that the CUL3 RNAi strategy targets less diverse cellular processes than the MLN4924 strategy as MLN4924 inhibits a upstream step of CRL activation [29] (see the KLHL12 RNAi section for a more specific strategy).

The increase in the levels of secreted and intracellular collagen suggests that collagen gene expression is increased by CUL3 depletion. To test this possibility, we measured levels of *COL1A1* mRNAs with a real time quantitative PCR (RT-qPCR) (Fig. 4E). As expected, we observed about 7-fold increase in the levels of COL1A1 mRNAs by CUL3 depletion. These results indicated that CUL3 regulates the expression of COL1A1.

### CUL3 is a regulator of the UPR of the ER

It was intriguing that the effect of CUL3 depletion on collagen became significant when the serum was present and that the effect of CUL3 depletion on collagen was lost when the serum was absent for 17h. Serum provides cells with growth factors that stimulate collagen synthesis [33]. It has been reported that a serum deprivation can induce ER stress (i.e., the UPR), which leads to inhibition of collagen synthesis in HSFs [34]. However, because we added back the serum after the transfection step and continued to incubate the cells in a serum containing medium for 48h, ER stress triggered by the serum withdrawal would be minimal. Nevertheless, we tested whether the serum withdrawal procedure could trigger ER stress, masking the effect of CUL3 depletion on collagen. For this purpose, we monitored changes in UPR signaling involving PERK, IRE1α, and ATF6α [35]. Remarkably, CUL3 depletion led to an increase in the levels of PERK and IRE1α, but not those of full-length (FL) ATF6α. (Fig. 4A, F, G). Note that when the UPR is activated ATF6α FL is transported from the ER to the Golgi where it is processed to the N-terminal fragment (NTF) [36]. Phosphorylation of eIF2α, a downstream molecule of the PERK signaling pathway [37], was enhanced by CUL3 depletion (Fig 4A, H). This enhancement became quite significant when the serum was present. However, when serum was absent, phosphorylation of eIF2α was only marginally enhanced by CUL3 depletion even though the levels of PERK were increased. Perhaps this was due to a low signal to noise ratio of p-eIF2α under the condition. However, it was clear that our serum withdrawal procedure did not trigger ER stress by itself (Fig. 4, compare lane 1 with lane 3 for PERK, IRE1α, p-eIF2α, and ATF6α). It should be noted that phosphorylation of eIF2α was enhanced by CUL3 depletion even in the absence of extrinsic ER stressors such as DTT, thapsigargin, and tunicamycin. These results suggest that CUL3 modulates levels of UPR sensors such as PERK and IRE1α and that CUL3 regulates PERK signaling.

Intrigued by this new CUL3-UPR connection, we asked whether the increase in the amounts of PERK and IRE1α leads to an overactivation of the UPR when cells are stressed with an extrinsic ER stressor. For this purpose, we treated HSFs with DTT (Fig. 5). DTT treatments retarded the mobility of PERK and IRE1α in SDS-PAGE (Fig. 5A). These shifts in mobility have been shown to be caused by phosphorylation [38]. The levels of PERK and IRE1α species including the slowly migrating ones were increased by CUL3 RNAi upon DTT treatments, but not those of ATF6α-FL, suggesting that UPR signaling branches involving PERK and IRE1α are upregulated (Fig. 5 A, C-E). We also monitored changes in downstream components of the UPR. Although phosphorylation of eIF2α was enhanced by CUL3 RNAi alone, the DTT treatments did not further enhance phosphorylation of eIF2α (Fig. 5A, F). XBP1 is a critical component of the IRE1α pathway and a UPR induction triggers synthesis of the spliced form of XBP1 [39, 40]. The DTT treatments induced synthesis of spliced XBP1 as expected (Fig. 5A). CUL3 RNAi combined with DTT treatments led to a significant increase of spliced XBP1 compared to the DTT treatments alone (Fig. 5A, G). CUL3 depletion did not change the levels of ATF6α-NTF (Fig. 5A, H). Based on these observations we concluded the CUL3 is also an important regulator of IRE1α signaling in HSFs.

**Figure 5.**
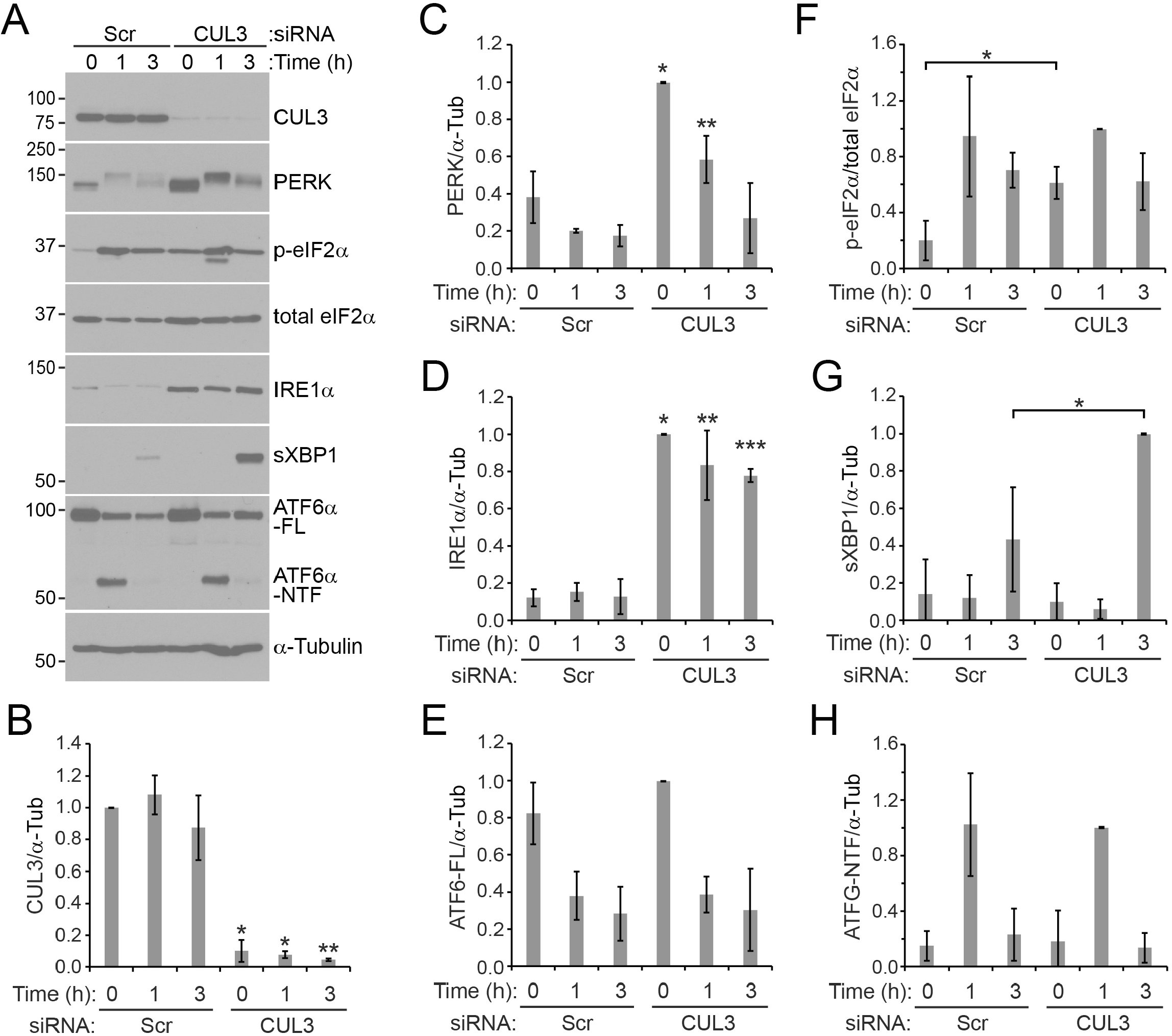
Regulation of IRE1α-signaling by CUL3. (A) HSFs were transfected with scrambled (Scr) or CUL3 siRNAs (50 nM) in the presence of the serum. Cells were treated with 2 mM DTT for an indicated duration. (A) A representative immunoblot images. Upon a DTT treatment, bands of PERK and IRE1α migrated slower than those in the untreated condition due to phosphorylation [38]. (B-H) Immunoblots images of 3 independent experiments were quantified. Statistical analyses were performed with Student’s t-test. Error bars represent standard deviations. (B) Comparison of CUL3 levels (normalized to α-tubulin) between presence and absence of DTT at each time point. P*<0.0001; P**<0.005, n=3. (C) Comparison of PERK levels between presence and absence of DTT at each time point. P*<0.005; P**<0.01, n=3. Note that band smearing prevented faithful quantification at 3h. (D) Comparison of IRE1α levels. P*<0.0001; P**<0.005; P***<0.0005, n=3. (E) Comparison of ATF6α levels. (F) Comparison of p-eIF2α levels normalized to total levels of eIF2α. P*<0.05, n=3. (G) Comparison of spliced (s) XBP1 levels. P*<0.05; n=3. (H) Comparison of ATF6α NTF levels.

### CUL3 regulates UPR sensors differentially in different cell lines

There are 183 BTB domain genes in the human genome which likely serve as a substrate-specific adaptor for CUL3 [5]. Different cell lines/tissues are expected to have different profiles of BTB-domain proteins. Thus, we asked if the CUL3-UPR link is observed in different cell types. To address this question we performed RNAi in IMR90 and HeLa (Fig. 6). The effects of CUL3 depletion on IRE1α and PERK were observed in IMR90 (Fig. 6 A and B). Interestingly, CUL3 depletion increased levels of PERK but not those of IRE1α in HeLa cells (Fig. 6A and C), indicating that CUL3 does not always regulate levels of PERK and IRE1α concurrently. HeLa cells do not express *COL1A1* (Fig. S4). Thus, the UPR dysregulation triggered by CUL3 depletion is not an indirect consequence of aberrant collagen export from the ER. Our results are consistent with the idea that CUL3 regulates the levels of PERK and IRE1α with different BTB domain proteins that are differentially expressed in HSFs, IMR90, and HeLa (Fig. 6D).

**Figure 6.**
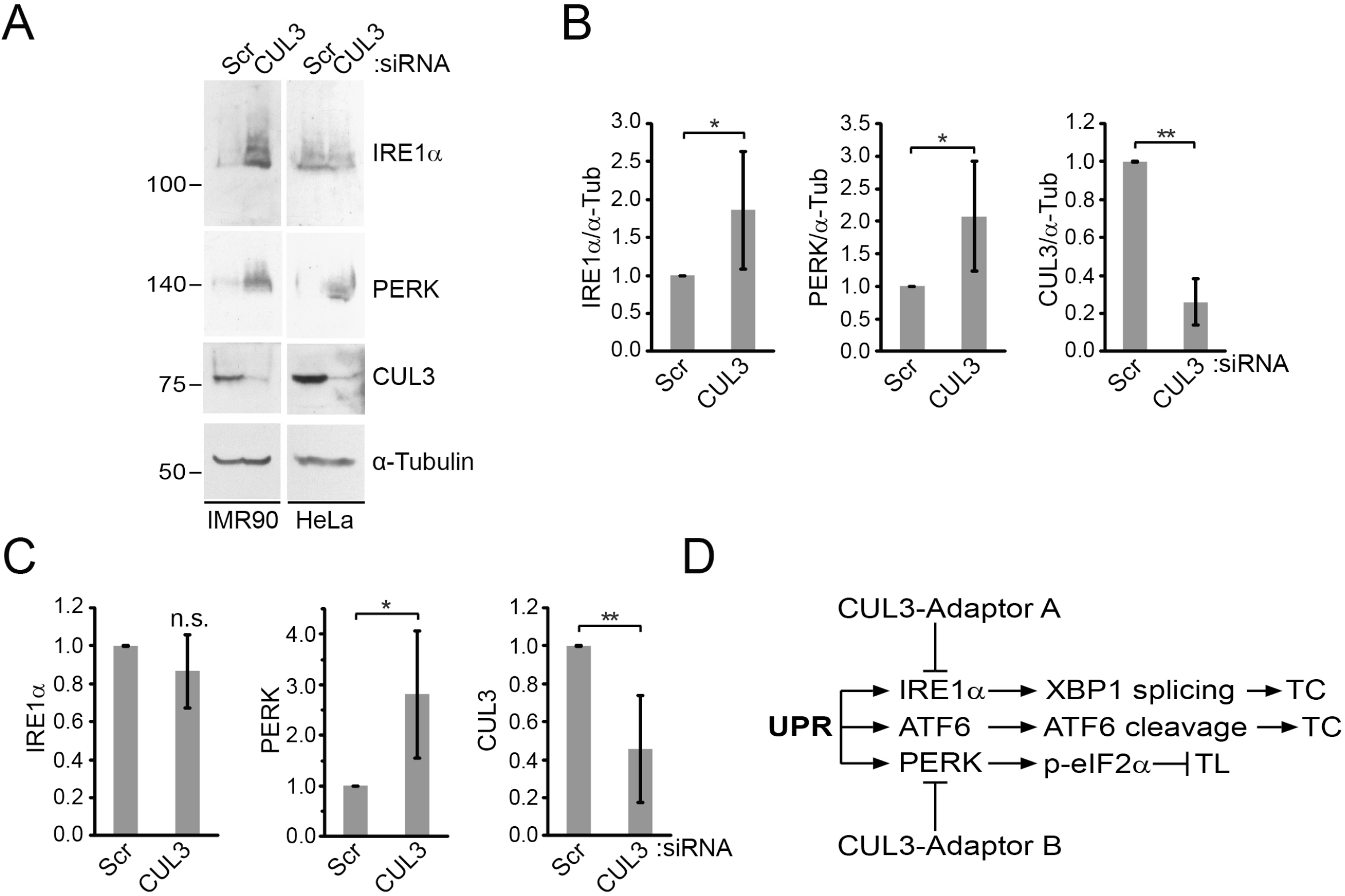
A tissue-specific regulation of the UPR by CUL3. (A, B, and C) Different cell lines were transfected with scrambled (Scr) or CUL3 siRNAs (50 nM) in the absence of the serum. Immunoblots were quantified for IMR90 cells (B) and for HeLa cells (C). Statistical analyses were performed with Student’s t-test. Error bars represent standard deviations. (B) P*<0.05, P**<0.0001, n=4. (C) P*<0.05; P**<0.005, n=4. (D) A model explaining the cell line-specific regulation of the UPR by CUL3 and its adaptors. TC, transcription; TL, translation.

### KLHL12 regulates collagen synthesis and the UPR

We then tested whether the effects of CUL3 on collagen and the UPR is mediated through KLHL12 in HSFs. KLHL12 depletion led to a reduction of both secreted and intracellular collagen (Fig. 7A-D), which coincided with a decrease of COL1A1 mRNA levels (Fig. 7G). These results suggest that KLHL12 regulates COL1A1 synthesis. In addition, KLHL12 depletion led to an increase in the levels of PERK and IRE1α when the serum was present during transfection, but not when the serum was absent during transfection (Fig. 7A, E, F). We believe this is partly due to inefficient KLHL12 depletion in the absence of the serum (Fig. 7B).

**Figure 7.**
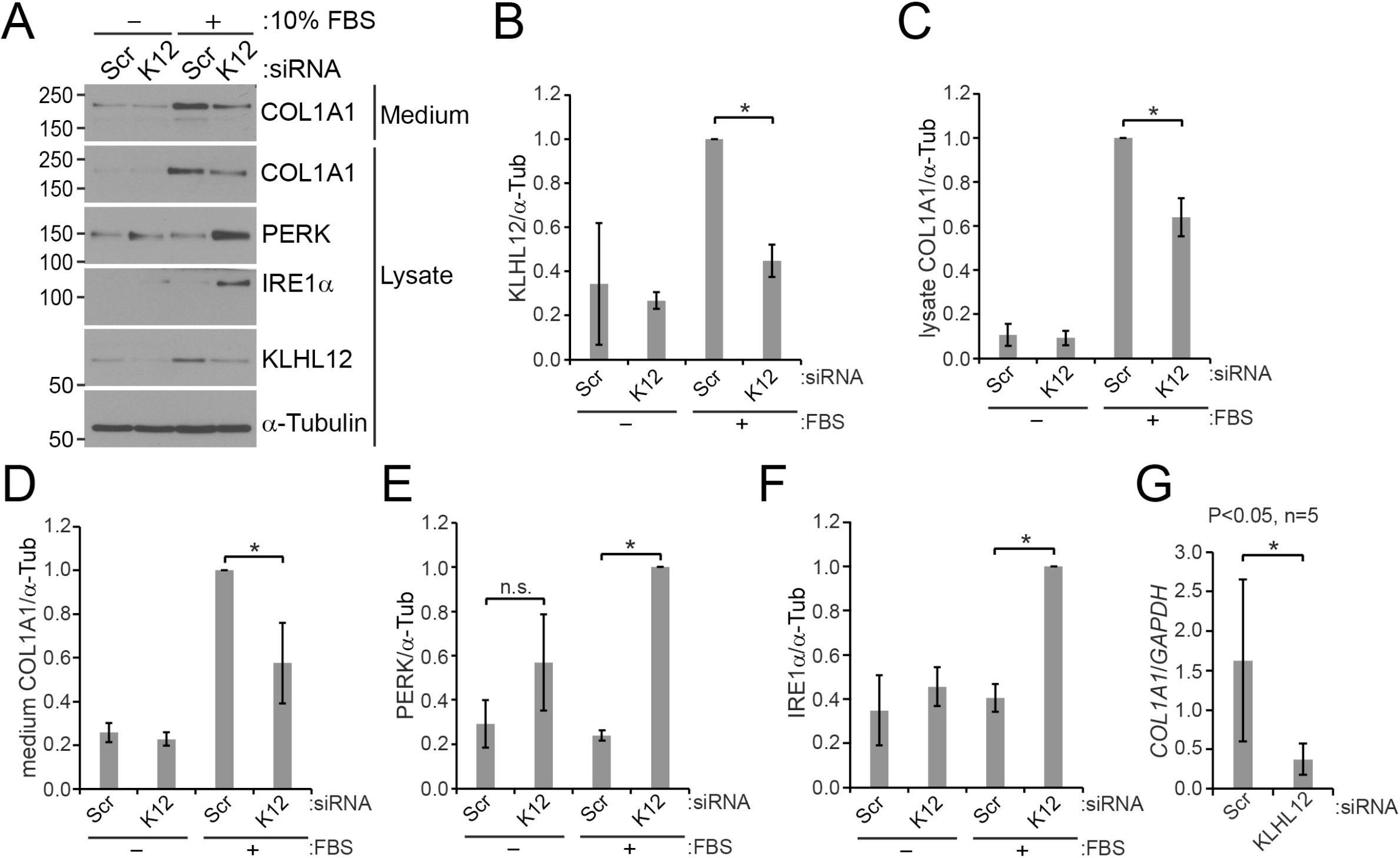
Regulation of COL1A1, PERK, and IRE1α levels by KLHL12. HSFs were transfected with scrambled (Scr) or KLHL12 (K12) siRNAs (50 nM) as described in the legend of Fig. 4. (A) A representative immunoblot images. (B-F) Immunoblots images of 3 independent experiments were quantified. Statistical analyses were performed with Student’s t-test. Error bars represent standard deviations. (B) Comparison of KLHL12 levels normalized to α-tubulin. P*<0.0005; n=3. Note that KLHL12 levels were increased in the presence of the serum for unknown reasons. (C) Comparison of intracellular COL1A1 levels. P*<0.005, n=3. (D) Comparison of secreted COL1A1 levels. P*<0.05, n=3. (E) Comparison of PERK levels. P*<0.0001, n=3. n.s., non-significant. (F) Comparison of IRE1α levels. P*<0.0001, n=3. (G) Levels of *COL1A1* mRNAs were measured with RT-qPCR and normalized to levels of *GAPDH* mRNAs. P*<0.05, n=5.

It is intriguing that CUL3 depletion and KLHL12 depletion resulted in opposite outcomes with respect to collagen synthesis, but similar outcomes with respect to the UPR sensor regulation. Possibly, this there are BTB-domain proteins that can regulate collagen synthesis positively or negatively and KLHL12 happens to be a positive regulator.

### A muscle-specific BTB domain protein regulate PERK levels

KLHL41 is a muscle-specific protein [41], the major BTB domain protein in muscle [42], and interacts with CUL3 [43]. We asked if KLHL41 can regulate the levels of the UPR sensors. To test this possibility we differentiated C2C12 myoblasts to myotubes (CMT) by incubating the cells in a differentiation medium containing 50 nM insulin (Fig. S6) [44]. Depletion of CUL3 or KLHL12 led to increased levels of PERK, but not IRE1α (Fig. 8). This result suggests the muscle-specific effect of CUL3 on PERK is conveyed through the muscle-specific BTB domain protein KLHL41. Surprisingly, when we performed similar experiments in the myotubes differentiated in the absence of insulin, we observe a reduction of PERK levels (Fig. S7). Apparently, the cells reacted differently to the depletion CRL3^KLHL41^, depending on the state they are in. This reflects a complexity in PERK regulation by CRL3^KLHL41^.

**Figure 8.**
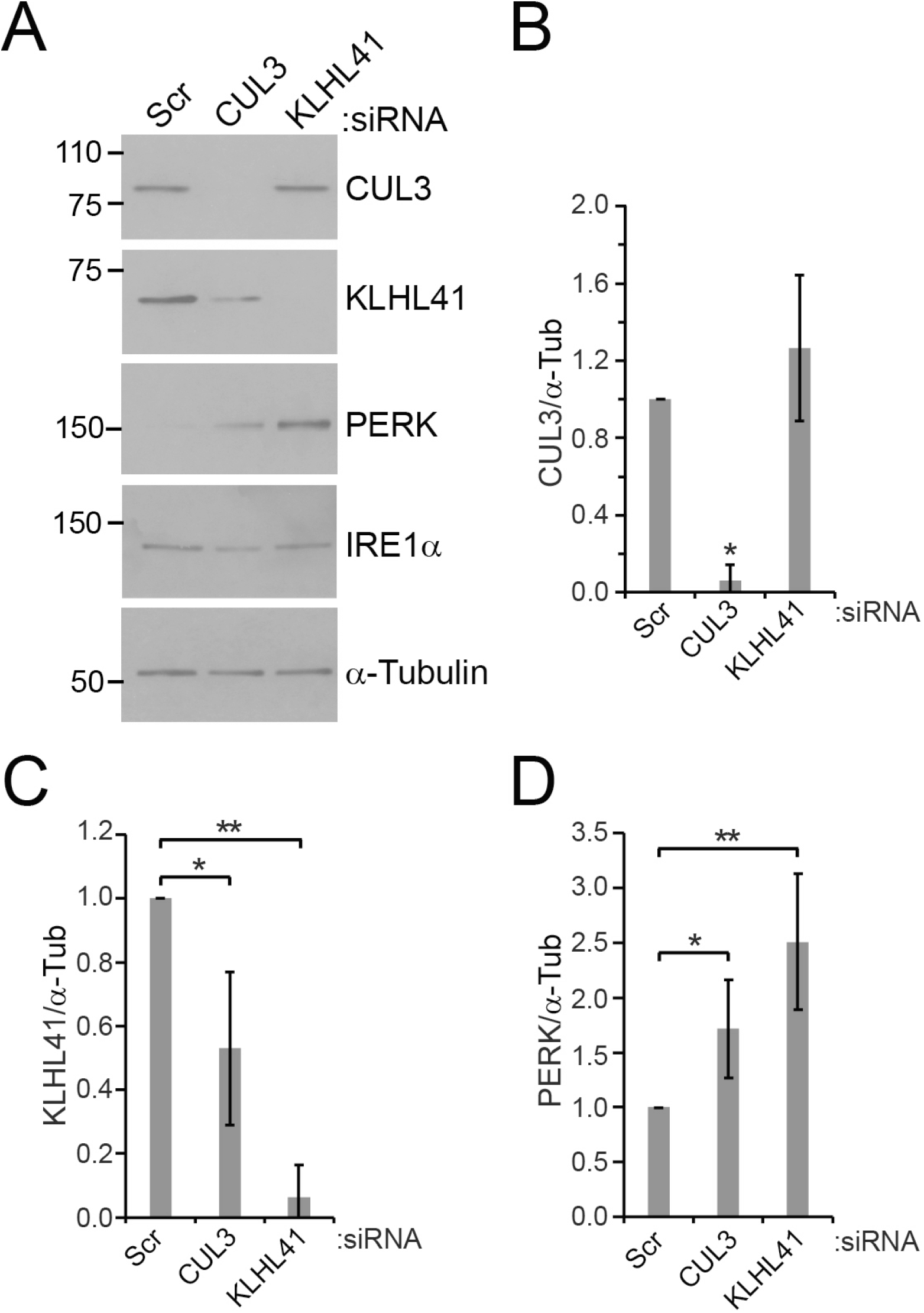
KLHL41 is responsible for PERK regulation in C2C12 mouse myotubes (CMTs). C2C12 mouse myoblasts were transfected with scrambled (Scr) or CUL3 or KLHL41 (K41) siRNAs (25 nM) in a differentiation medium in the presence of insulin (50 nM) at day 0. The medium was replaced with a fresh differentiation medium containing insulin at day 1. Cells were harvested at day 3 or 4. (A) A representative immunoblot images. For unknown reasons, a depletion of CUL3 led to a decrease of KLHL41 levels, but not vice versa. (B-D) Quantifications of 5 independent experiments were plotted. Statistical analyses were performed with Student’s t-test. Error bars represent standard deviations. (B) Comparison of CUL3 levels (normalized to α-tubulin). P*<0.0001, n=5. (C) Comparison of KLHL41 levels. P*<0.005, P**<0.0001, n=5. (D) Comparison of PERK levels. P*<0.01, P**<0.001, n=5.

## DISCUSSION

Our data support the idea that the integrity of CRL3^KLHL12^ is critical for formation of KLHL12-COPII-coated structures as MLN4924 treatments or KLHL12 mutants compromised KLHL12-COPII structures. Surprisingly, depletion of CUL3 or KLHL12 affected collagen levels in the cell as well as in the culture medium when the siRNA transfection was performed in the presence of the serum in HSFs. However, similar depletion did not influence secretion, nor intracellular levels of collagen when the siRNA transfection was performed in the absence of the serum. Although the physiological relevance of these two *in vitro* conditions remains to be determined, a quantitative proteomic study has revealed that levels of COL2A1 and COL9A1 are enhanced in skeletal muscles of skeletal muscle-specific *CUL3* knockout mice [45]. The changes of the collagen levels in our experiments appear to be linked to collagen expression as the transcripts of COLA1 were affected. Based on these results we conclude that CUL3 and KLHL12 play a role in regulating collagen synthesis. As we did not observe the effect of CUL3 on collagen secretion there seems to exist a CRL3-independent collagen secretion pathway. Thus, it is unlikely that CRL3^KLHL12^ is essential for collagen export in every cell types and tissues. Instead, different cell types or tissues seem to have alternative or compensatory mechanisms for exporting collagen from the ER.

One of the salient aspects of our findings is that CRL3^KLHL12^ regulates the UPR. Because of the increased levels of PERK and IRE1α due to CRL3 depletion, downstream UPR signaling was activated accordingly in HSFs, indicating that CRL3 regulates UPR signaling. Although the effect of CUL3 on collagen synthesis was influenced by the serum, the effect of CUL3 on the UPR sensors was not affected by the serum. These results suggest that the CUL3-UPR interaction is independent of any possible effects of serum withdrawal. In addition, the CUL3-UPR interaction does not depend on COL1A1 expression because the CUL3-PERK interaction was observed in HeLa cells where *COL1A1* is not expressed.

If CUL3 depletion had triggered gross ER stress, we would have observed an activation of all three signaling pathways of the UPR. However, ATF6α levels and its proteolytic processing were not affected by CUL3 depletion (Fig. 5A, compare lane 2 with lane 5 for ATF6α). Thus, we do not think CUL3 depletion induces gross ER stress. In addition, ER stressors such as DTT, thapsigargin, and tunicamycin do not alter the levels of the UPR sensors, but rather modify them (phosphorylation or proteolytic processing) (Fig. 5A, compare lane 1 with lane 2 for PERK, IRE1α, and ATF6α). Our results are consistent with the idea that CUL3 influences the UPR in a specific fashion (i.e., PERK and IRE1α). This specific effect likely involves a specific BTB domain protein(s). Our data indicate that KLHL12 is one of them. In addition, although we have not found a cell line where CUL3 regulates ATF6, it is possible that CUL3 regulates ATF6 in a certain cell line through a BTB-domain protein.

KEAP1, a BTB-domain protein, regulates UPR signaling involving NRF2 [46]. NRF2 is a transcription factor which regulates the expression of antioxidant genes in response to oxidative stress [47, 48]. NRF2 levels are maintained via association with KEAP1 [49]. This association is regulated by PERK-dependent phosphorylation of NRF2 [50]. This mode of UPR regulation by CRL3^KEAP1^ is distinct from the newly identified mode of UPR regulation by CRL3^KLHL12^ where KLHL12 regulates the levels of PERK and IRE1α. Thus, CUL3 and its adaptor molecules influence the UPR with multi-faceted strategies, emphasizing fundamental importance of CRL3s in UPR regulation.

We discovered that the regulation the UPR by CUL3 varies in different cell lines and culture conditions: 1) CUL3 depletion increased levels of PERK and IRE1α in HSF and IMR90, 2) CUL3 depletion increased levels of PERK, but not those of IRE1α in HeLa cells and CMTs, and 3) CUL3 depletion affected PERK oppositely in CMTs depending on differentiation conditions. We believe that these differential effects arise from differentially expressed BTB-domain proteins and their substrate(s). CUL3 recognizes its substrate molecules through a BTB-domain protein [6]. There are 183 BTB-domain genes in the human genome [5]. They are likely expressed differentially in various tissues. In fact, KLHL41 is expressed predominantly in muscles [51]. The cellular phenotypes induced by depletion of KLHL41 were similar to those observed with depletion of CUL3 in CMTs, suggesting that the effect of CUL3 is relayed through KLHL41. In addition, a BTB-domain protein can recognize several different target molecules, vastly expanding functional roles of CUL3 [43]. Importance of tissue-specific expression of a substrate molecule(s) of KLHL41 was also obvious in our data. Mere absence of KLHL41 is not sufficient to lower the levels of PERK. This is because PERK levels are relatively high in cell lines (i.e., HSFs) which do not express KLHL41. In addition, although KLHL41 is expressed much less in myoblasts than in myotubes, the levels of PERK are quite high in myoblasts compared to those in myotubes (Fig. 6S, compare day 0 to day 2-5). Thus, a myotube-specific substrate(s) of KLHL41 likely plays a critical role in PERK regulation. The identity of the substrate remains to be determined.

## ACKNOWLEDGEMENTS

We would like to thank the laboratories of Michael Rape and Randy Schekman for sharing valuable resources with us. Research reported in this publication was supported by the National Institute of General Medical Sciences of the National Institutes of Health under award number R01GM110373.

## AUTHOR CONTRIBUTION

KK, TM, SP, JK designed the study; KK, TM, SP collected the data; KK, JK wrote the article and all contributed to editing and reviewing the submission.

## The abbreviations

BTB: Bric a brac, Tramtrack and Broad-Complex
CMT: C2C12 mouse myotube
CRL: CUL3-RING ubiquitin ligase
ECM: extracellular matrix
ER: endoplasmic reticulum
FBS: fetal bovine serum
HSF: human skin fibroblast
ned: neddylated
RNAi: RNA interference
siRNA: small interfering RNAs
Ub: ubiquitin
UPR: unfolded protein response

