## Supplemental Information for "Cullin3-RING ubiquitin ligases are intimately linked to the unfolded protein response of the endoplasmic reticulum"

### ABSTRACT

CUL3-RING ubiquitin ligases (CRL3s) are involved in various cellular processes through different Bric a brac, Tramtrack and Broad-Complex (BTB)-domain proteins. KLHL12, a BTB-domain protein, appears to play an essential role in export of large cargo molecules like procollagen from the endoplasmic reticulum (ER). It has been suggested that $CRL3^{KLHL12}$ mono-ubiquitinates SEC31 and mono-ubiquitinated SEC31 increases the dimension of a COPII coat to accommodate the large cargo molecules. As we examined this model, we found that functional $CRL3^{KLHL12}$ was indeed critical for the assembly of large COPII structures. Interestingly, we noticed that $CRL3^{KLHL12}$ influences collagen synthesis in human skin fibroblasts (HSFs). Our results also suggest that there is a $CRL3^{KLHL12}$–independent collagen secretion route in HSFs. In addition, we found that $CRL3^{KLHL12}$ strongly influences levels of sensors of the unfolded protein response (UPR). Different cell lines reacted differently to CUL3 depletion with respect to UPR regulation. This cell line-dependency appears to rely on a cell line-specific BTB-domain protein(s). Consistent with this idea, depletion of a muscle-specific BTB-domain protein KLHL41 recapitulated the effects of CUL3 depletion in C2C12 myotubes in UPR regulation. Based on these results we propose that $CRL3^{KLHL12}$ and $CRL3^{KLHL41}$ are regulators of the UPR.

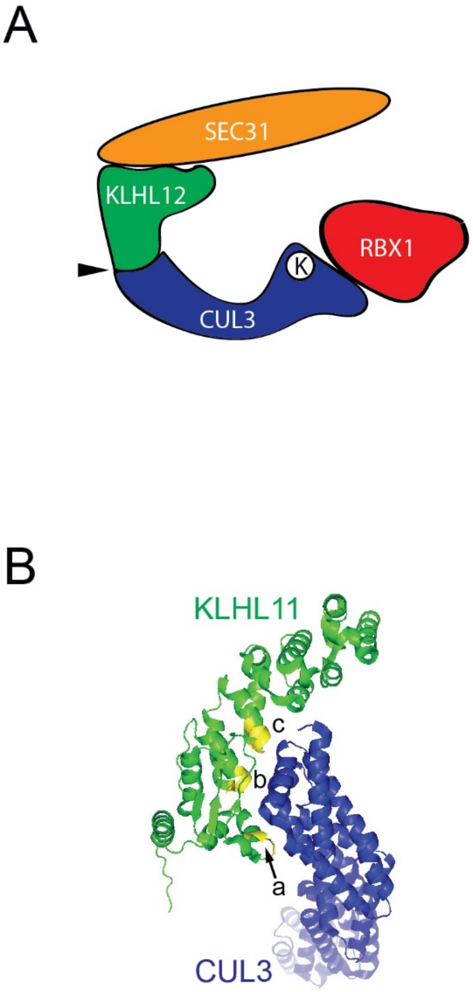


C

```
                         a
 54 CSDYFCAMFTSELSEKGKPYVDIQ   KLH12
118 ATEYFTPLLSGQFSESRSGRVEMR   KLH11
 73 ASDYFKAMFTGGMKEQDLMCIKLH   KLHL9
103 SSPVFKAMFTNGLREQGMEVVSIE   KEAP1
 71 CSPYFCAMFTGDMSESKAKKIEIK   KLHL3
 77 CSPYFHAMFTGEMSESRAKRVRIK   KLHL2
221 RSPVFSAMFEHEMEESKKNRVEIN   SPOP
 89 CSPYFRAMFTGELAESRQTEVVIR   KLH20

                          b
 99 TVENVQELLPAACLLQLKGVKQAC   KLH12
169 STGSVHEVLELADRFLLIRLKEFC   KLH11
118 NMDNLQDTLEAASFLQILPVLDFC   KLHL9
148 GEKCVLHVMNGAVMYQIDSVVRAC   KEAP1
116 TEENVQVLLPAASLLQLMDVRQNC   KLHL3
122 TEENVQVLLPAAGLLQLQDVKKTC   KLHL2
266 LDKMADDLLAAADKYALERLKVMC   SPOP
134 EEGNVQTLLPAACLLQLAEIQEAC   KLH20

                       c
134 NCLGIRDFAETHNCVDLMQAAEVF   KLH12
204 NCVAIHSLAHMYTLSQLALKAADM   KLH11
153 NCVEVGRIANTYNLIEVDKYVNNF   KLHL9
183 NAIGIANFAEQIGCVELHQRAREY   KEAP1
151 NCLGIRAFADVHTCTDLLQQANAY   KLHL3
157 NCLGIRAFADMHACTDLLNKANTY   KLHL2
301 NAAEILILADLHSADQLKTQAVDF   SPOP
169 NCLGIRAFADTHSCRELLRIADKF   KLH20
```

Figure S1. Mutations of KLHL12 in the KLHL12-CUL3 interface. (A) A schematic illustration of a CRL$^{KLHL12}$ with SEC31. KLHL12 mutations were introduced to putative CUL3 binding sites (arrowhead). (B) An X-ray crystallographic structure of the KLHL11-CUL3 complex (4APF) [1]. KLHL11 and KLHL12 are 27% identical. Mutations were introduced to

KLHL12 based on 4APF and amino acid sequence homology (C). Mutation sites (a, b and c) were marked in yellow. (C) Sequence comparison of the selected human BTB-Kelch proteins near the mutation sites. Highly conserved residues are in boldface. The mutated residues are underlined. The selected residues were replaced with alanines.

Table S1. Primers used to generate KLHL12 Mut A, Mut B, and Mut C using pcDNA5/FRT/TO KLHL12-FLAG

| Mutation | Primer | Sequence |
|---|---|---|
| KLHL12 Mut A (HindIII/StuI) | Kel F (966) | GCGTTTAAACTTAAGCTTGGTACCGAGCTCGGATCCatgggaggc |
| | Kel R (1223) | aacataaggtttccccttAGCTGCGGCAGCactagtgaacatggcacagaagt |
| | Kel F (1171) | acttctgtgccatgttcactagtGCTGCCGCAGCTaaggggaaaccttatgtt |
| | Kel R (1796) | CCGAAGTTCAGGCCTCAGATGAAACTTCTT |
| KLHL12 Mut B (HindIII/StuI) | Kel F (966) | GCGTTTAAACTTAAGCTTGGTACCGAGCTCGGATCCatgggaggc |
| | Kel R (1367) | gcaggcttgtttcacaccTGCAGCGGCTGCGGCAGCggctgcaggaagcagttc |
| | Kel F (1314) | gaactgcttcctgcagccGCTGCCGCAGCCGCTGCAggtgtgaaacaagcctgc |
| | Kel R (1796) | CCGAAGTTCAGGCCTCAGATGAAACTTCTT |
| KLHL12 Mut C (HindIII/StuI) | Kel F (966) | GCGTTTAAACTTAAGCTTGGTACCGAGCTCGGATCCatgggaggc |
| | Kel R (1456) | tgcttgcatcaggtcaacAGCGGCAGCTGCGGCTGCAGCatccctaatacccaggca |
| | Kel F (1406) | tgcctgggtattagggatGCTGCAGCCGCAGCTGCCGCTgttgacctgatgcaagca |
| | Kel R (1796) | CCGAAGTTCAGGCCTCAGATGAAACTTCTT |

We performed a series of PCR reactions using pcDNA5/FRT/TO KLHL12-FLAG as a template and the DNA oligomers shown above as primers to generate KLHL12 mutant constructs. For Mut A, a PCR reaction was performed with primers Kel F (996) and Kel R (1223) to synthesize 258 bp fragments and with primers Kel F (1171) and Kel R (1796) to synthesize 626 bp fragments. These two fragments were purified and then used as templates to synthesize 831bp-fragments with primers Kel F (996) and Kel R (1796). The 831 fragments were digested with HindIII and StuI and inserted into the corresponding sites of pcDNA5/FRT/TO KLHL12-FLAG. For Mut B, a PCR reaction was performed with primers Kel F (996) and Kel R (1376) to synthesize 402 bp fragments and with primers Kel F (1314) and Kel R (1796) to synthesize 483 bp fragments. These two fragments were purified and then used as templates to synthesize 831bp-fragments with primers Kel F (996) and Kel R (1796). The 831 fragments were digested with HindIII and StuI and inserted into the corresponding sites of pcDNA5/FRT/TO KLHL12-

FLAG. For Mut C, a PCR reaction was performed with primers Kel F (996) and Kel R (1456) to synthesize 495 bp fragments and with primers Kel F (1406) and Kel R (1796) to synthesize 393 bp fragments. These two fragments were purified and then used as templates to synthesize 831bp-fragments with primers Kel F (996) and Kel R (1796). The 831 fragments were digested with HindIII and StuI and inserted into the corresponding sites of pcDNA5/FRT/TO KLHL12-FLAG.

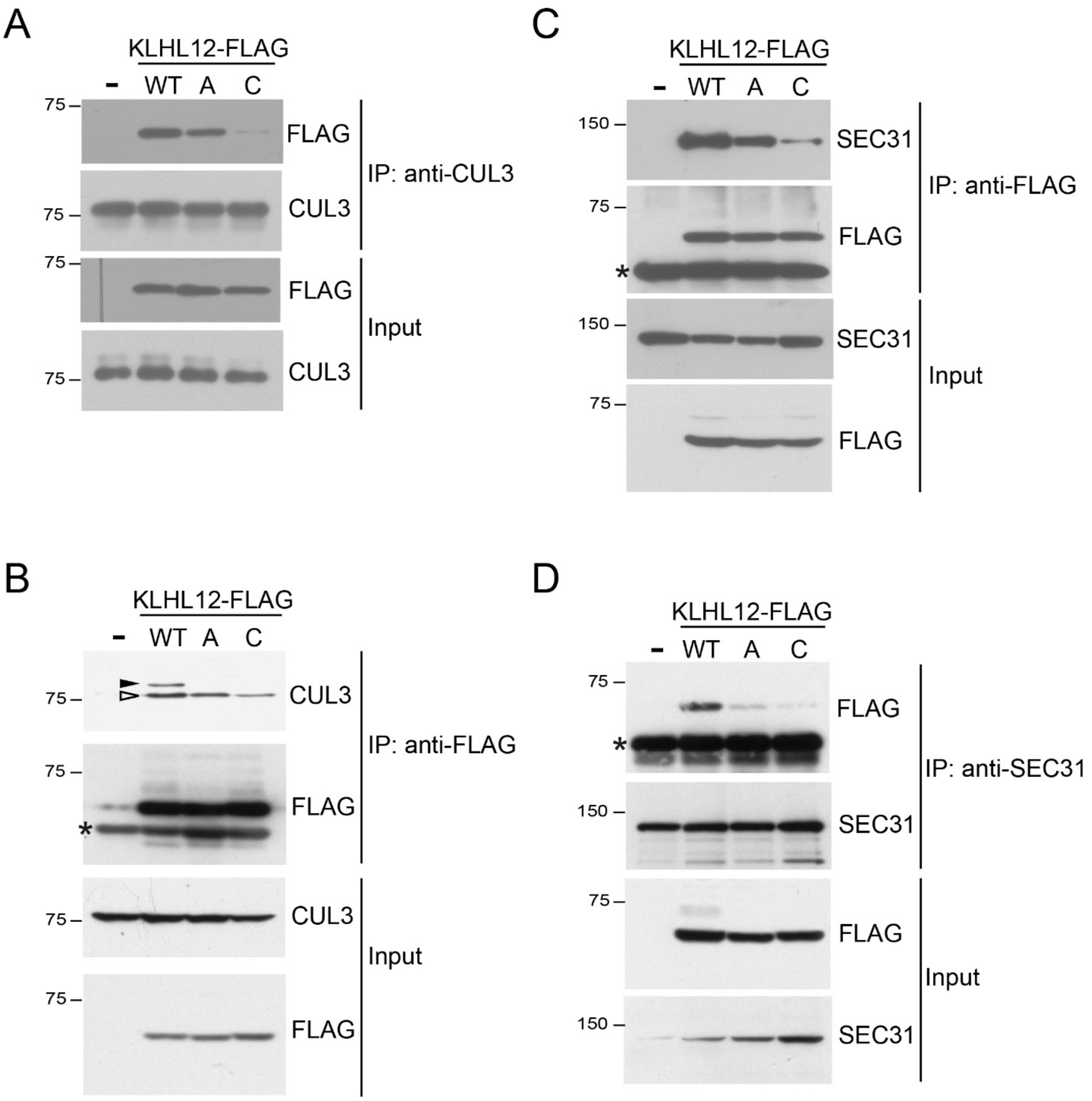


Figure S2. The KLHL12-CUL3 interaction influences the KLHL12-SEC31 interaction. (A-D) Stable 293 cells harboring an empty vector (-) or an indicated KLHL12-FLAG construct were treated with doxycycline to induce the expression of a KLHL12-FLAG construct. Cellular lysates were processed for immunoprecipitation using an indicated antibody. The closed arrowhead and the open arrowhead represent neddylated and unneddylated CUL3, respectively. *Immunoglobulin. WT, wildtype; A, mutant A; C, mutant C.

For coimmunoprecipiation, cells were scraped and collected by centrifugation. The collected cells were lysed by douncing 30 times using lysis buffer (0.1% NP-40, 2.6 mM KCl, 1.5 mM $KH_2PO_4$, 140 mM NaCl, 8 mM $Na_2HPO_4$-$7H_2O$, and 1X protease inhibitor cocktail) on ice. After centrifugation for 15 min at 14,000 rpm, cleared lysates were incubated with indicated antibodies overnight at 4°C, followed by incubation with protein A or protein G Sepharose beads (GE Healthcare, Milwaukee, WI) for 3 h at 4°C with gentle rocking. The beads were harvested by centrifugation and washed three times with the lysis buffer. After adding the SDS-sample buffer and heating for 5 min at 95 °C, proteins were resolved with SDA-PAGE, transferred to PVDF membrane (IPVH00010, Millipore, Bedford, MA, USA), and probed with primary antibodies and subsequently with horseradish peroxidase-conjugated secondary antibodies.

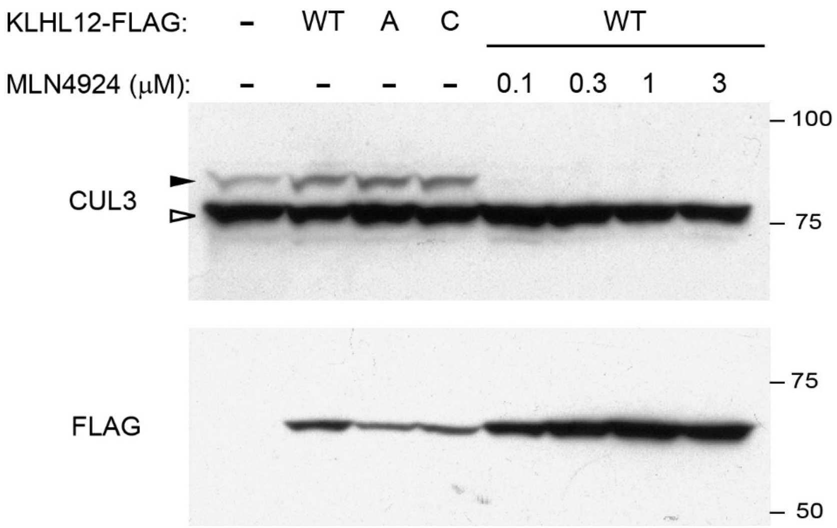


Figure S3. Neddylation of CUL3 occurs normally when mutant KLHL12 constructs are expressed. HEK cells stably expressing KLHL12-FLAG constructs were incubated in the presence or absence of MLN4924. Expression of KLHL12-FLAG was induced with doxycycline. The filled arrowhead and the open arrowhead represent neddylated and unneddylated CUL3, respectively. An MLN4924 treatment inhibited neddylation of CUL3 and stabilized KLHL-FLAG.

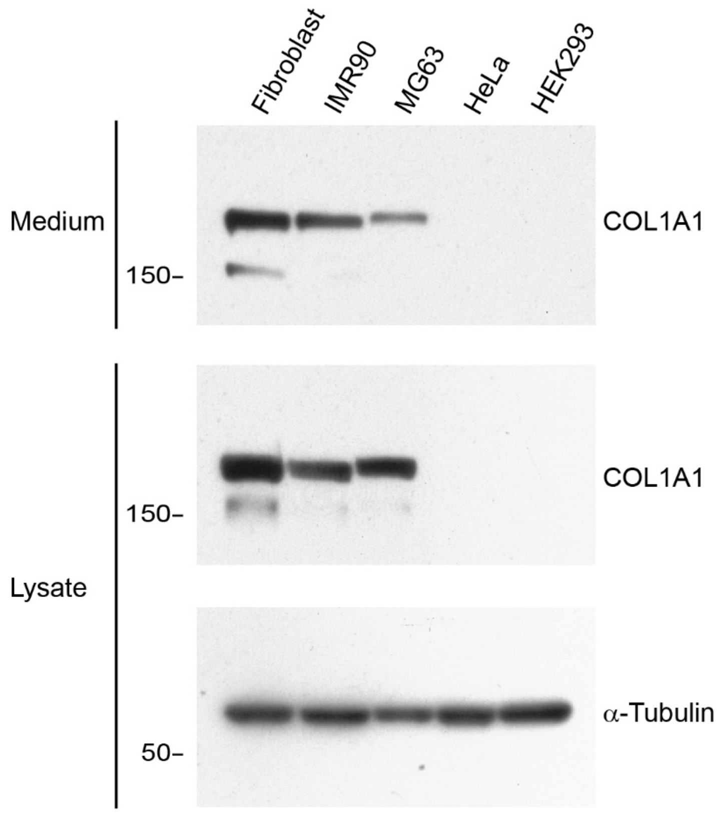


Figure S4. Expression of *COL1A1* in different cell lines. Confluent cells were incubated with fresh medium and incubated for 24h. Cells and conditioned media were collected and processed for immunoblotting.

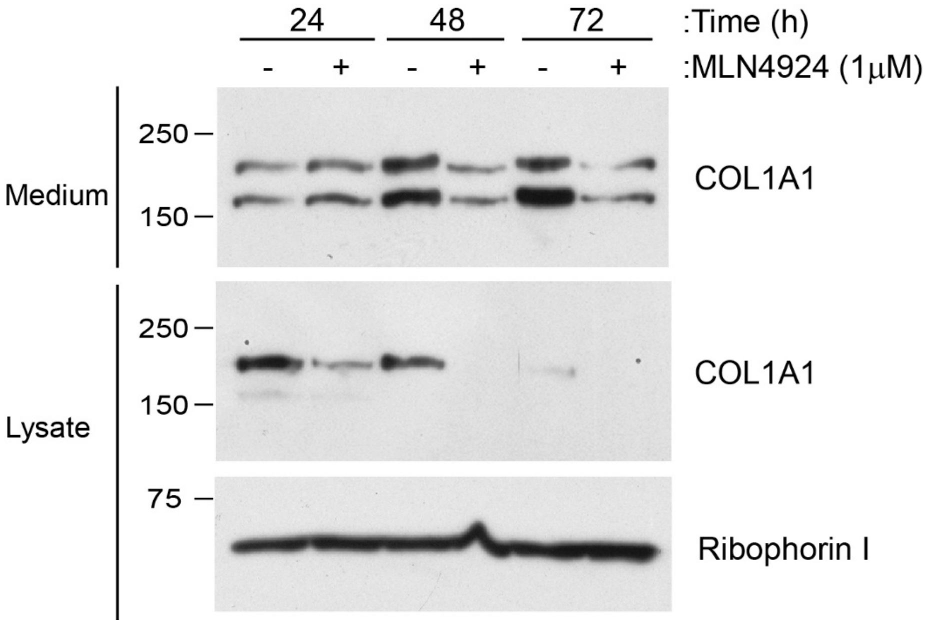


Figure S5. Regulation of cellular COL1A1 levels by MLN4924. HSFs were plated and then incubated for 24h. The culture medium was replaced with fresh medium with or without MLN4924 (1μM). Conditioned media and cells were collected at indicated times and processed for immunoblotting. The amount of a medium loaded into the gel was normalized to the amount of total proteins in a cell lysate. Riboporin I was probed as a loading control.

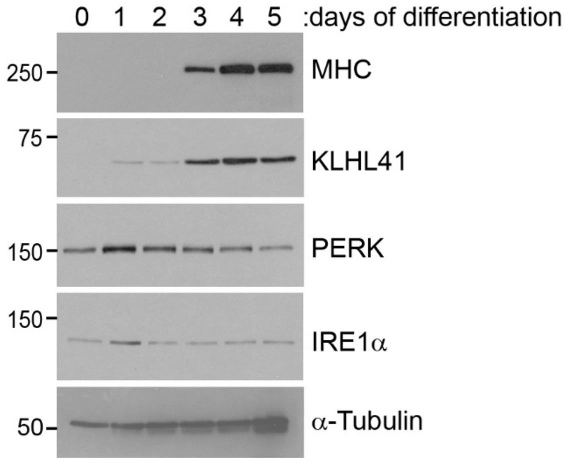


Figure S6. Differentiation of C2C12 myoblasts to myotubes. C2C12 myoblasts were 50 nM insulin. C2C12 myoblasts were differentiated in a differentiation medium containing 50 nM insulin [2]. Muscle markers such as KLHL41 and myosin heavy chain (MHC) were robustly expressed as differentiation progressed [3].

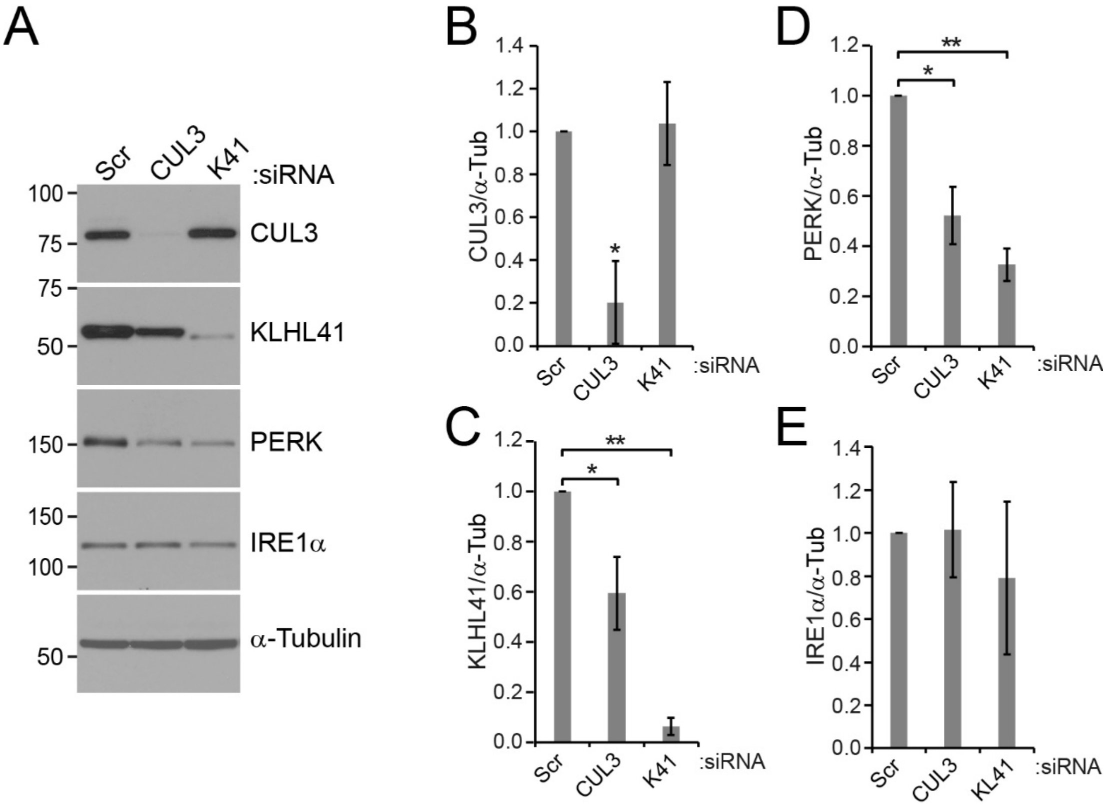


Figure S7. KLHL41 regulates PERK in C2C12 myotubes. C2C12 mouse myoblasts were transfected with scrambled (Scr) or CUL3 or KLHL41 (K41) siRNAs (25 nM) in a differentiation medium at day 0. The medium was replaced with a fresh differentiation medium at day 1. Cells were harvested at day 3 or 4. (A) A representative immunoblot images. For unknown reasons, a depletion of CUL3 led to a reduction of KLHL41 levels, but not vice versa. (B-E) Quantifications of 3 independent experiments were plotted. Statistical analyses were performed with Student's t-test. Error bars represent standard deviations. (B) Comparison of CUL3 levels (normalized to α-tubulin). P*<0.005, n=3. (C) Comparison of KLHL41 levels. P*<0.01, P**<0.0001, n=3. (D) Comparison of PERK levels. P*<0.005, P**<0.0001, n=3. (E) Comparison of IRE1a levels (normalized to α-tubulin). Not significant, n=3.
